# Profound impact of hippocampal output on the interpretation of tactile input patterns in SI neurons

**DOI:** 10.1101/2022.11.27.518101

**Authors:** Leila Etemadi, Jonas M.D. Enander, Henrik Jörntell

## Abstract

Due to continuous state variations in neocortical circuits, individual SI neurons *in vivo* display a variety of intracellular response types to repeated presentations of the exact same tactile input pattern. The specific intracellular response obtained depends on a time-evolving combination of the specific input with the current neocortical state. To manipulate the internal cortical state, we here used brief electrical stimulation of the output region of the hippocampus, which preceded the delivery of specific tactile afferent input patterns to digit 2 of the anesthetized rat. We find that hippocampal output had a diversified and remarkably strong impact on the specific set of intracellular response types each SI neuron displays to each given tactile input pattern. The findings show that hippocampal output can profoundly impact the state-dependent interpretation of tactile inputs in SI neurons and hence influence their perception, potentially with affective and semantic components.

## Introduction

Analysis of the human brain has indicated that the neocortex is functionally a heavily interconnected network (Bullmore and Sporns, 2009). We have previously shown that the processing of tactile information also at the cellular level in SI is dependent on other parts of the cortex. This was shown both by stroke-like lesions and by electrical manipulation of brain activity in neocortical areas with a remote location to the SI (Etemadi et al., 2022; Wahlbom et al., 2019). Recent anatomical studies have indicated that the extent of connections made by individual corticocortical axons have previously been greatly underestimated (Ahrlund-Richter et al., 2019; Gerfen et al., 2018) (see also https://www.janelia.org/project-team/mouselight; https://mouse.braindatacenter.cn/), hence giving structural support for these observations. Given that perception to some extent depends on expectation, which through the internal cortical state may exert impact already before the tactile afferent information enters the SI cortex, these findings suggest that tactile perception could be influenced by a globally integrated action across the neocortex.

The hippocampal formation is closely interconnected with the neocortex. The hippocampus receives sensory information from a large number of different neocortical areas via the entorhinal cortex into the subiculum (Schultz and Engelhardt, 2014). Indeed, the hippocampus is also engaged in processing of tactile input (Pereira et al., 2007). Through its output via the subiculum, the hippocampus can monosynaptically activate the anterior thalamus (Winnubst et al., 2019) and indirectly thereby also widespread areas of the cortex. Subiculum neurons can also have widespread projections, such as to the hypothalamus, regions of the temporal/entorhinal cortices surrounding the hippocampus, and the ventral striatum (Matsumoto et al., 2019; Winnubst et al., 2019), which also enables the hippocampus to potentially impact the activity across widespread areas of the neocortex.

Using intracellular recordings from SI neurons, we have previously shown that repeated presentations of a given spatiotemporal tactile input pattern from digit 2 combine with the state of the neocortical network to result in a variety of response states in SI neurons (Norrlid et al., 2021). The relative orthogonality of these response states suggested the presence of attractor-type dynamics (Norrlid et al., 2021; Ringach, 2009), reminiscent of what has also been proposed to characterize the neuron population activity dynamics in the hippocampus (Gardner et al., 2022; Gardner et al., 2019). More recently, we showed that manipulation of the internal cortical state through electrical activation of a remote neocortical area can change the preferred network solutions, or the types of response states observed in SI neurons, for every given tactile input pattern (Etemadi et al., 2022). Remarkably, this change occurred despite that the remote cortical perturbation did not induce any overt response in the recorded neurons. That finding suggested that the effects on the response states were indirectly mediated via more subtle changes in the state of the neocortex globally, which in turn would alter the number of ‘open’ network pathways supplying the recorded neuron with information related to the tactile event. Such gating of input pathways and information distribution across the neocortex is likely part of the central mechanisms for generating perception, and alteration of such pathways could be a root cause of illusions. Here, we address the question if also the hippocampal output, generated by electrical activation of its neural output stage in the subiculum, can impact the responses in SI neurons to given spatiotemporal tactile input patterns. We find that the hippocampal activation dramatically altered the responses of the SI neurons to tactile inputs, suggesting that the state of hippocampal activity could have a profound effect on the cortical interpretation, or perception, of such inputs.

## Results

We made whole cell patch clamp recordings from putative pyramidal neurons (see Methods; also (Norrlid et al., 2021)) between layers III-V in the forepaw region of the SI cortex, while stimulating the distal part of the second digit with eight fixed spatiotemporal tactile activation (TA) patterns delivered in pseudorandom order (Figure 1A), a protocol we’ve used in numerous previous publications (Enander and Jorntell, 2019; Enander et al., 2019; Etemadi et al., 2022; Norrlid et al., 2021; Oddo et al., 2017; Wahlbom et al., 2019). Here, these TA input patterns were provided in isolation or in combination with a preceding brief stimulation of the subiculum/terminal CA1 region of the hippocampus (HIPTA) (Figure 1A-C). As the study relied on comparisons between individual raw responses with large variance, we needed a high number of repetitions of each stimulus, and we could therefore include only recordings which maintained a high quality for a duration of at least 25 minutes (N=9 neurons).

**Figure 1.**
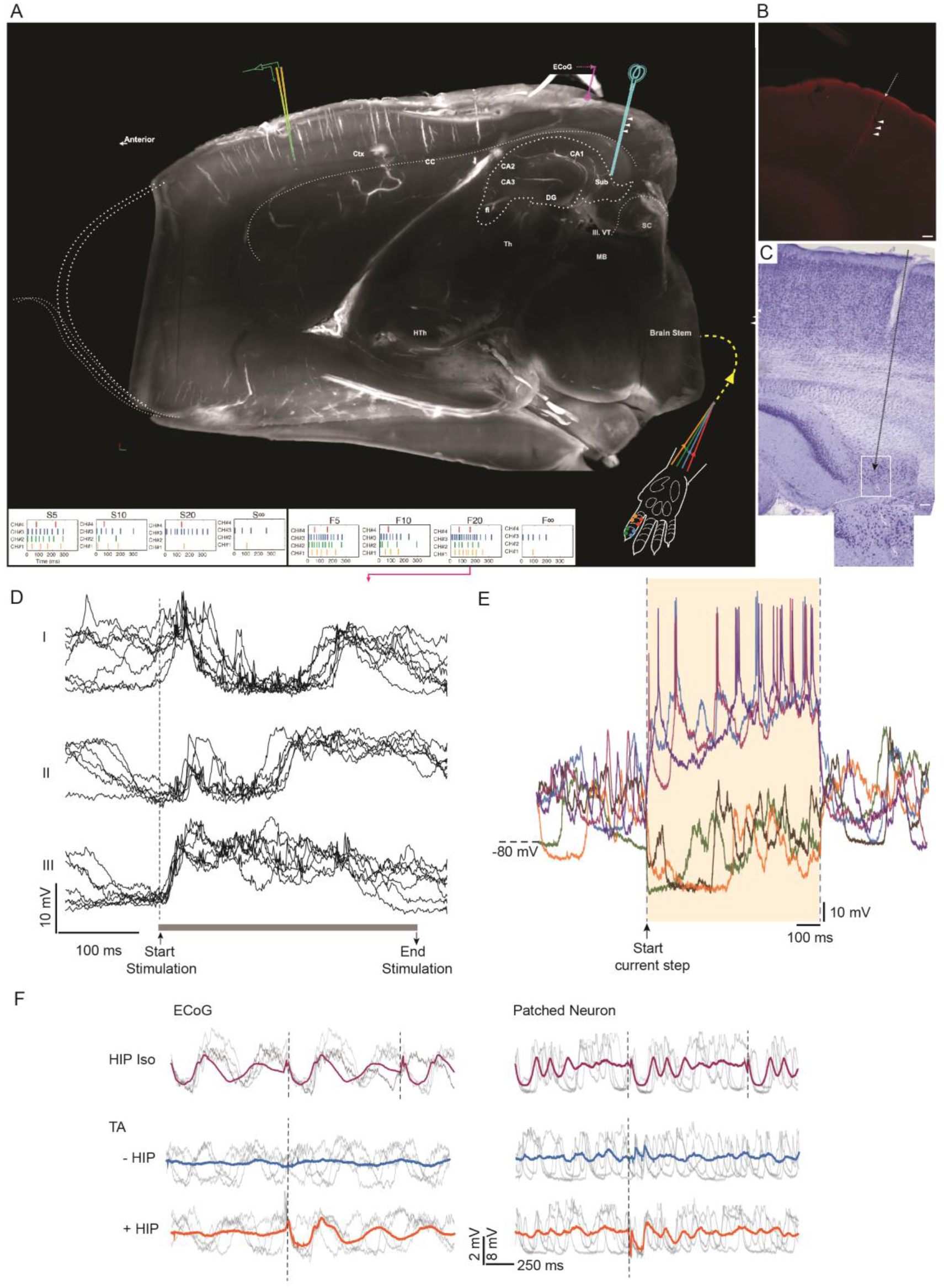
Experimental setup. (A) Location of the stimulation and recording electrodes visualized in one hemisphere clarified via the iDISCO Clearing protocol and visualized using light-sheet microscopy. White arrowheads indicate identified location of the HIP stimulation electrode (blue cartoon electrode, with the tip located in the subiculum/terminal CA1 region). In the bottom part of this image, the second digit of the left forepaw along with the location of four pairs of electrodes (color coded, corresponding to different tactile ‘channels’, ‘CH’) are shown together with the eight different spatiotemporal patterns of tactile stimulation (TA). The patch-clamp recording (indicated by a yellow/green cartoon electrode) was made in the cortical SI area. (B) HIP stimulation electrode track identified within the clarified brain tissue (white arrowheads). The stimulation electrode was covered in neurobiotin solution before the experiment. The presented image is a snapshot captured by light-sheet microscopy following conjugating neurobiotin with the streptavidin Alexa Flour-568 (red color in the image). Scale bar: 0.1 mm. (C) Histological identification of a HIP stimulation electrode track in regular light microscopy using cresyl violet acetate staining technique. The location of the tip of the HIP electrode in the subiculum area is visible in the white square area as a slight ‘burn’ mark, shown also with further magnification below. Scale bar: 0.1 mm. (D) Examples of intracellular response types (I-III) evoked by repeated stimulations using the tactile input pattern F20. The dashed line indicates the starting point and the grey line indicates the duration of the stimulation. Some remnants of the blanked shock artefacts created minor spikes in the recording. (E) Intracellular responses evoked by current step commands (+0.6 nA and -0.6 nA, respectively) used for controlling the recording quality. (F) ECoG (left) and intracellular responses (right) to the isolated subiculum stimulation (HIP iso), the TA input pattern F∞ without and with preceding HIP stimulation (HIPTA). Thick traces in each subpanel represent averages. Grey thin traces are example raw data traces. Dashed vertical lines indicate the onsets of the stimulations (which occurred at a higher frequency for the HIP iso condition).

As we have reported earlier (Etemadi et al., 2022; Norrlid et al., 2021), each individual neuron in SI cortex displayed a wide variety of response types when the exact same TA stimulus was repeatedly presented (Figure 1D). This indicates that response averaging would risk missing essential features of these cortical responses, as averaging would tend to overemphasize the impact of the peripheral component of the response versus its internally generated components. Since the latter presumably is essential to the perceptual process in the cortex, which we were interested in here, we instead focused on the characterization of the pool of individual raw responses evoked by each stimulus pattern and each stimulus condition. This process involved the step to cluster the responses evoked by the same stimulus pattern. We used a clustering method based on the temporal profile of the evoked responses (Figure 1D; see also (Etemadi et al., 2022)), where the temporal response profile reflects the temporal structure of the responses in the different subnetworks forwarding input to the recorded neuron (Etemadi et al., 2022). Clustering relied on analysis of spike-free intracellular responses which reflect the synaptic input to the cell. In each recording we therefore applied a mild hyperpolarization current to prevent the neuron from spiking (the occasional few remaining spikes could be blanked in software), whereas the quality of the recording was occasionally verified by injecting depolarizing current to evoke spike responses (Figure 1E).

Stimulation of the subiculum (HIP) in isolation (iso) typically evoked detectable responses in the average ECoG signal (recorded from the parietal cortex, Figure 1A), which were several order of magnitudes longer than the duration of the stimulation (7 pulses at 333 Hz, i.e. 18 ms), whereas the TA stimulation alone evoked no detectable response (Figure 1F, left column). In the average intracellular signal of the recorded SI neurons, this HIP response was often much less distinct and typically less long lasting than in the ECoG (Figure 1F, right column). In contrast, the average response to the TA stimulus was always clear in the intracellular recordings from the SI neurons (though widely different from on neuron to the other, see (Norrlid et al., 2021)). Notably, the intracellular responses evoked by HIP iso and by the TA pattern, respectively, were clearly different from the HIPTA response, indicating that the effect on the global cortical state induced by the HIP stimulation was non-linearly combined with the impact of the TA input to produce a new response.

We found that the combination of a TA input with the preceding HIP stimulation could generate different outcomes across different TA input patterns. Figure 2 illustrates examples of the impacts the HIP stimulation could have on the average responses evoked by four different TA patterns in the same cell. Notably, the hippocampal output caused the S10 TA input pattern to completely lose an early excitatory component, whereas a later excitatory component was instead greatly amplified. For the S5 TA input pattern, the early excitatory component was in contrast not eliminated, in fact a bit amplified but also slightly slowed down. Also the impacts on the two other example TA input patterns F5 and F10 were highly different.

**Figure 2.**
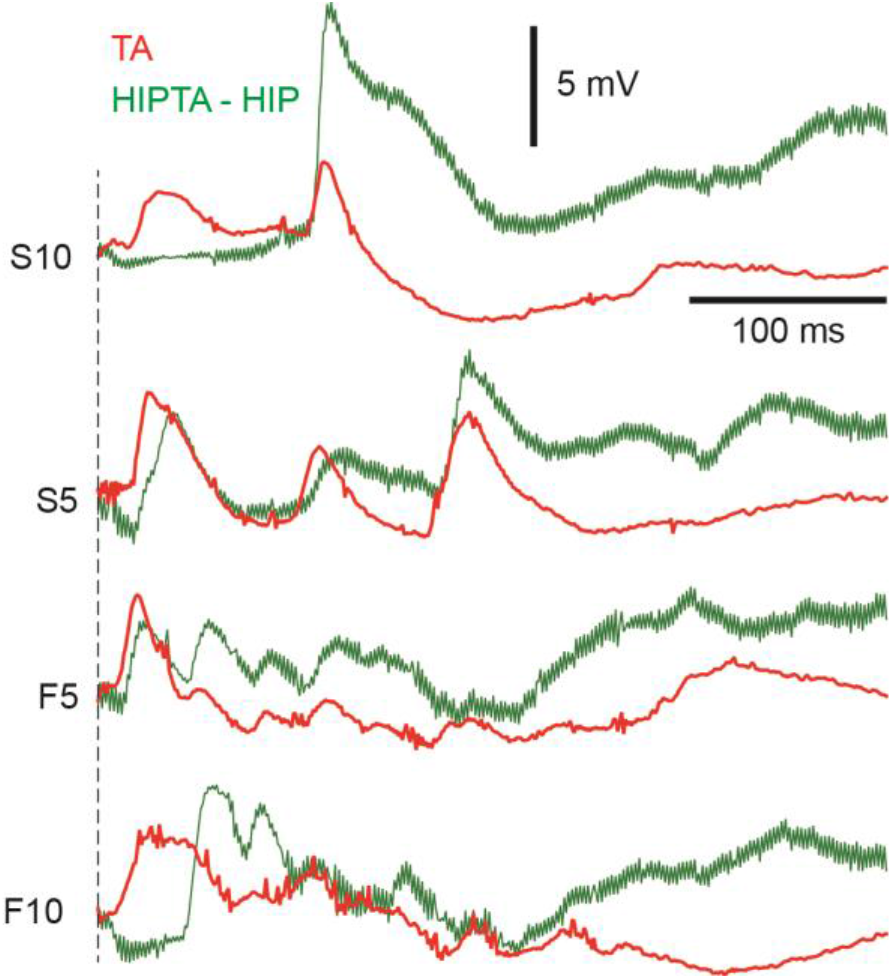
Impact of hippocampal stimulation (HIP) on average responses evoked by specific TA patterns. The red traces illustrate the average intracellular response to the indicated TA input. The green traces are the corresponding average traces evoked by the same TA inputs when they were combined with the HIP stimulation (HIPTA). The average response to the isolated HIP stimulation was subtracted from the HIPTA responses to illustrate only the TA component of the HIPTA responses (HIPTA-HIP).

### Profound impact of the HIP stimulation on the TA response types

We next made a detailed analysis of the response types evoked in the SI neurons. Different intracellular response types evoked by the same TA input pattern is a striking phenomenon that we have previously reported (Etemadi et al., 2022; Norrlid et al., 2021). Response types identified among a set of individual raw responses can be demonstrated with response clustering, and here we explored if these response type clusters formed by the pool of raw responses evoked by a single TA input pattern were impacted by preceding HIP stimulation. Figure 3A illustrates some example response types evoked by the F5 TA input. Figure 3B presents responses to the same TA input pattern but now preceded by the brief HIP stimulation. In this case, the response types evoked by the TA input pattern were profoundly altered compared to Figure 3A. This can for example be seen in that response peaks were eliminated or added at different points in time during the duration of the TA input pattern when it was combined with preceding HIP stimulation.

**Figure 3.**
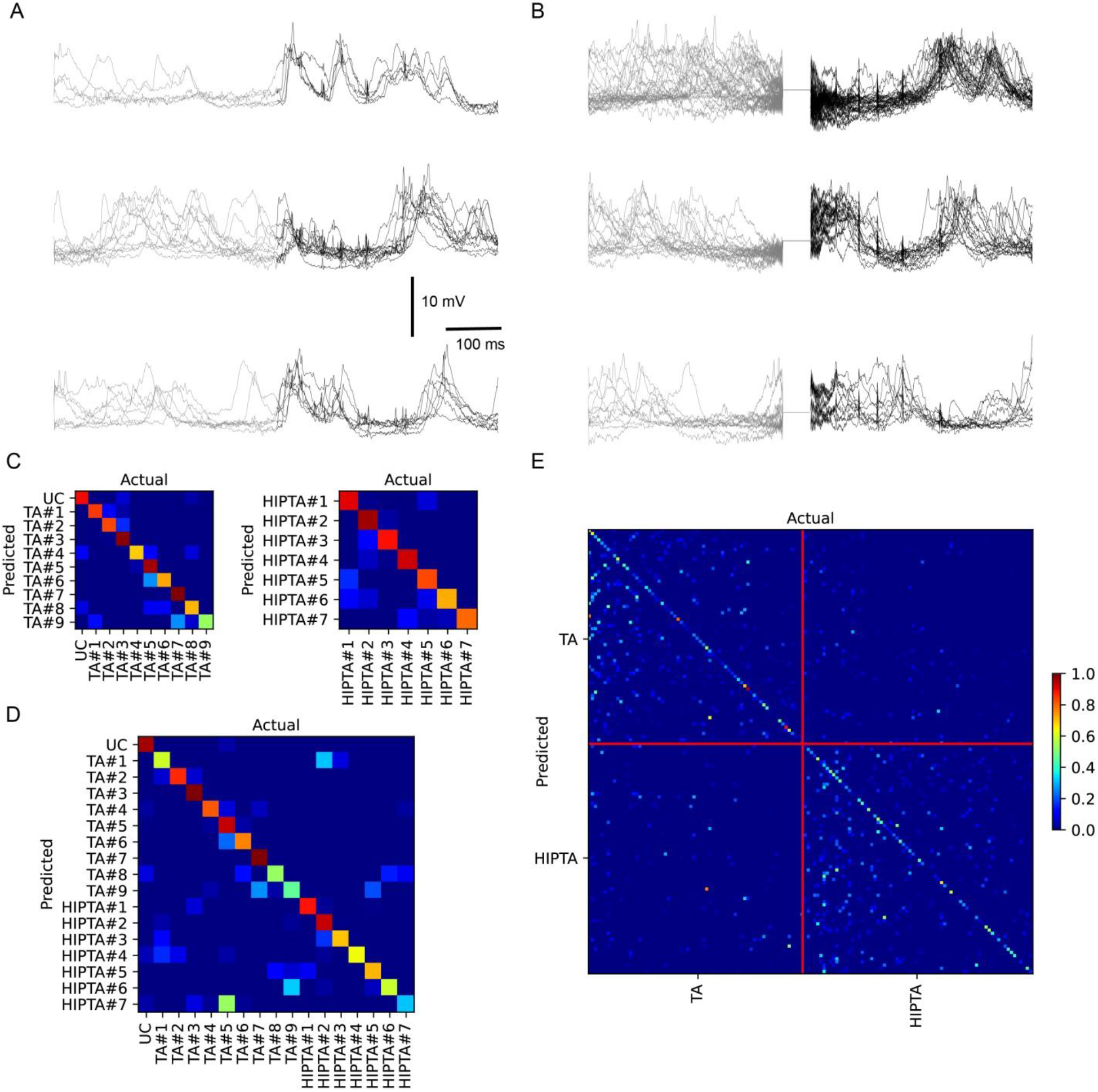
The intracellular response types evoked by a given tactile input pattern (TA) were profoundly altered when combined with subiculum stimulation (HIPTA). (A) Examples of superimposed raw intracellular responses for three response types evoked by repeated activation of one specific tactile input pattern (F5). Black part of the traces represent the time period when there was ongoing TA input, the preceding grey part of the traces were recorded before the onset of the TA input pattern. (B) Response types evoked in the same cell and using the same tactile input pattern when preceded by the HIP stimulation. The HIP stimulation occurred during the blanked period of the recording. Note that some blanked shock artefacts of the TA input pattern left some remnants, which almost looked like small spikes in the intracellular signals. (C) Confusion matrix for the nine response types evoked by the F5 tactile input pattern, following a mapping of these response types to principal component space. The presence of a ‘non-blue’ diagonal in the matrix indicates that the identified response types, evoked by one and the same stimulation pattern, were highly distinct from each other (as previously analyzed in detail in Norrlid et al 2021; Etemadi et al 2021). UC, unclassified responses. (D) Similar display for the seven identified response types when the F5 tactile input pattern was preceded by hippocampal output. (E) Confusion matrix for the response types evoked by the F5 pattern under the two conditions (TA and HIPTA, without and with preceding hippocampal output). (F) Confusion matrix for the responses evoked by all eight tactile input patterns (Fig 1A) across the two conditions TA and HIPTA. Red vertical and horizontal lines divide the two conditions.

In order to verify that the differences in response types observed in the above example occurred systematically, we performed a quantitative comparison of the temporal profiles of the responses using a PCA+kNN classification analysis (see Methods; same approach as that used in (Etemadi et al., 2022)). We first compared the clusters evoked by the same TA input pattern, but separated based on the condition (i.e. with or without preceding HIP stimulation). This analysis showed that the nine clusters of response types evoked by this TA input pattern were systematically distinct, with little confusion (similarity) between the response types (Figure 3C). This was also true for the seven clusters evoked by the F5 HIPTA input condition (Figure 3D). But the more critical comparison was that between the TA and the HIPTA clusters. Figure 3E shows that all response types, except the TA#5 and the HIPTA#7 response clusters, were well separated from each other.

Whereas the preceding panels only illustrated responses evoked by the F5 example TA input, Figure 2F illustrates that even when all response clusters for all eight stimulation patterns, across the two input conditions, were compared, the confusion between all of these response clusters was still highly limited. These results hence indicate that the HIP conditioning stimulus profoundly impacted the response types evoked by each given TA input, and also confirmed our previous findings that the different response types evoked by each TA pattern is relatively unique compared to the response types evoked by other TA input patterns (Norrlid et al., 2021).

Across the population of neurons we recorded from, these quantitative measures were highly consistent (Table 1). We also observed that the responses that each given TA input pattern evoked varied greatly between the different neurons (not shown), which we have previously reported more systematically (Enander et al., 2019; Norrlid et al., 2021; Oddo et al., 2017).

**Table 1.** Separability of the clusters for each clustering and classification context as measured by the mean F1-score (extracted from confusion matrices of the types shown in Figure 3). The data in this table is the mean and standard deviation from the entire population of recorded cells. Note that the chance level depends on the number of clusters and varies for each neuron and pattern and cell, and that all F1 scores were many multiples above chance level. The HIP suffix refers to TA patterns preceded by HIP stimulation. The ‘Pooled’ suffix refers to when response clusters from the TA and HIPTA was compared together. The ‘Cell’ context represents classification across all TA and HIPTA clusters for a neuron.

| Figure | Context | Mean F1 (±s.d.) | Mean chance level (±s.d.) | N (cells) |
| --- | --- | --- | --- | --- |
| 3C | F5 | 83% (±7%) | 14% (±3%) | 9 |
| - | S5 | 82% (±5%) | 17% (±4%) | 9 |
| - | F10 | 83% (±6%) | 15% (±3%) | 9 |
| - | S10 | 80% (±6%) | 15% (±3%) | 9 |
| - | F20 | 83% (±7%) | 15% (±4%) | 9 |
| - | S20 | 80% (±4%) | 14% (±4%) | 9 |
| - | F∞ | 83% (±4%) | 15% (±7%) | 9 |
| - | S∞ | 83% (±7%) | 17% (±8%) | 9 |
| 3D | F5 HIP | 82% (±7%) | 16% (±3%) | 9 |
| - | S5 HIP | 80% (±7%) | 14% (±3%) | 9 |
| - | F10 HIP | 79% (±6%) | 15% (±5%) | 9 |
| - | S10 HIP | 83% (±7%) | 16% (±5%) | 9 |
| - | F20 HIP | 83% (±7%) | 15% (±3%) | 9 |
| - | S20 HIP | 79% (±8%) | 16% (±5%) | 9 |
| - | F∞ HIP | 82% (±8%) | 15% (±2%) | 9 |
| - | S∞ HIP | 81% (±7%) | 17% (±4%) | 9 |
| 3E | F5 pooled | 66% (±15%) | 7% (±1%) | 9 |
| 3E | S5 pooled | 67% (±14%) | 8% (±2%) | 9 |
| 3E | F10 pooled | 66% (±12%) | 7% (±2%) | 9 |
| 3E | S10 pooled | 69% (±8%) | 8% (±2%) | 9 |
| 3E | F20 pooled | 68% (±11%) | 7% (±1%) | 9 |
| 3E | S20 pooled | 66% (±11%) | 7% (±2%) | 9 |
| 3E | F∞ pooled | 65% (±12%) | 7% (±2%) | 9 |
| 3E | S∞ pooled | 68% (±10%) | 8% (±2%) | 9 |
| 3F | Cell | 23% (±11%) | 0.9% (±0.1%) | 9 |

### Hippocampal conditioning generated unique response clusters in SI neurons

Whereas the analysis in Figure 3 assumed that the TA and the HIPTA response types were different sets, and therefore clustered independently of each other, Figure 4 instead assumes that TA and HIPTA responses were part of the same distribution. Therefore, in this case the clusters were identified without an a priori separation of the responses based on condition. Importantly, the clustering algorithm could still identify clusters which with a very high probability consisted of responses of only one or the other condition. This is illustrated for one example neuron, for two different TA input patterns, F5 and S20, in Figure 4A,B. Across all eight TA input patterns, for the same example neuron, this was a consistent finding (Figure 4C). In our population of recorded neurons (N=9), almost all tended to a bimodal distribution (Figure 4D), whereas only two of the neurons had a more Gaussian-like distribution (for example the neuron represented by brown bars in Figure 4D). Hence, these findings underscore that the HIP conditioning stimulation profoundly impacted the response types evoked by each given TA input, since the clusters tended to contain responses from only one or the other condition.

**Figure 4.**
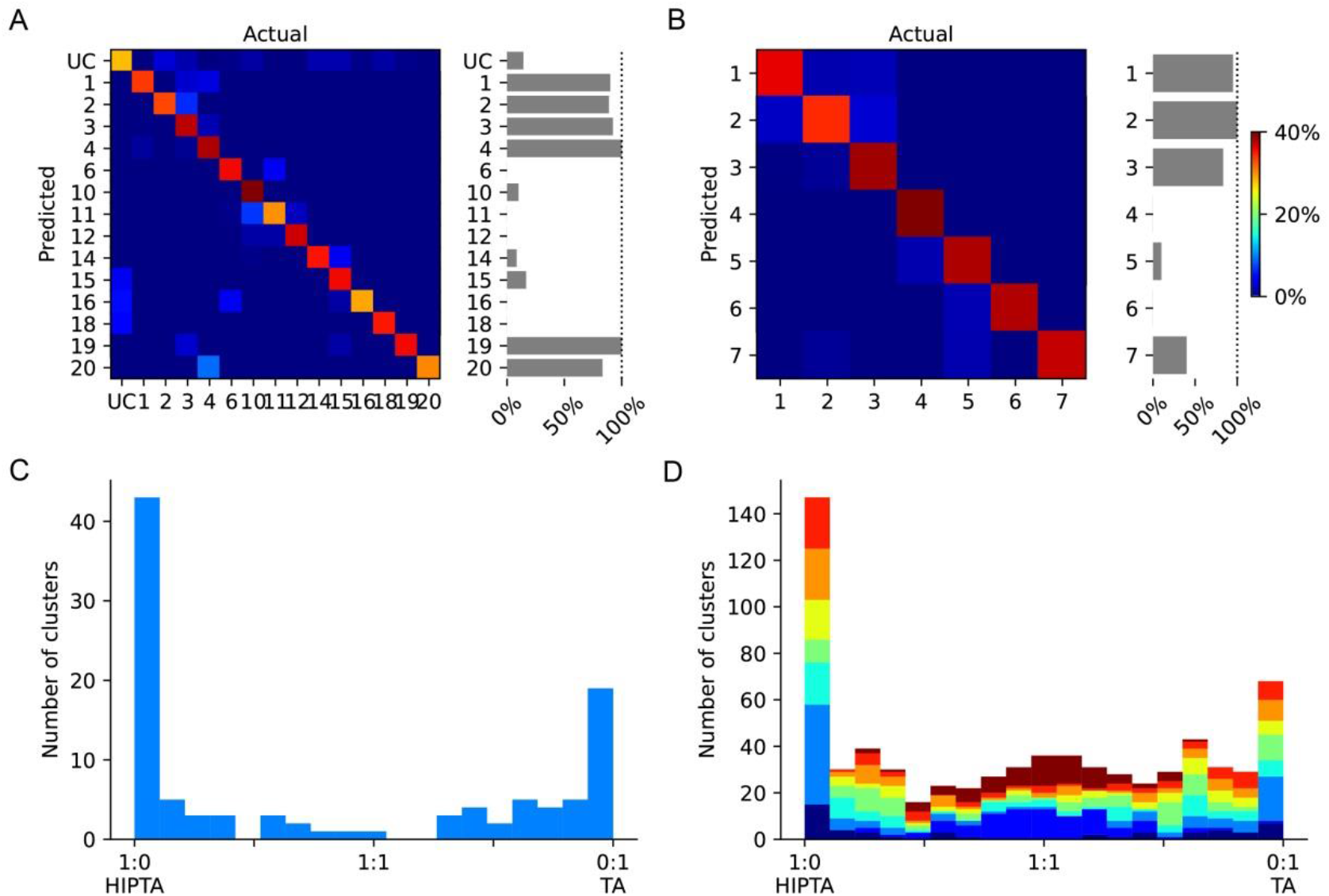
When the clustering algorithm was agnostic of the condition, identified response types were still relatively unique for each condition. (A) Confusion matrix for all response types evoked by one tactile input pattern (F5, the TA and the HIPTA conditions pooled), and the percentage of responses evoked with preceding hippocampal output for each response type. UC, unclassified responses. (B) Similar display for the responses evoked by another tactile input pattern (S20). (C) For each response type identified in this cell, across all TA input patterns, the proportion of the two conditions under which they were evoked. (D) Similar display as in (C) but for all cells recorded. Each cell was represented by a unique color.

## Discussion

Here we showed that brief preceding electrical activation of the subiculum, an input-output hub of the hippocampus, profoundly impacted the responses to tactile input patterns in SI neurons, located a long distance from the subiculum/CA1 terminal region (Figure 1A). As we have previously described (Etemadi et al., 2022; Norrlid et al., 2021), the variety of intracellular response states evoked by each individual tactile input pattern indicates that there is a range of dynamically gated cortical subnetworks supplying the sensory input to each neuron. The cortical network can thereby be described as containing a wide range of network solutions for any given spatiotemporal pattern of sensory input. What we described here is that the specific cortical network solution that applies at a given moment also depends on hippocampal activity. Hence, hippocampus is part of what defines the potential circuitry mechanisms underlying perception, where the internal state of the cortical circuitry would be equivalent to cortical ‘expectation’ or ‘prediction’.

Though the relative physical distance between SI and subiculum is large, neurophysiologically the distance is not so large. The subiculum projects monosynaptically to the anterior thalamus. From the thalamus, the subiculum output may reach the SI cortex within just a few synapses. There are also other potential pathways, for example to hypothalamus and ventral striatum and then to thalamus, which is another potential route by which hippocampal output can impact SI activity. Even so, the magnitude of the impact that we observed here was surprising. The synchronized activation that was achieved by the hippocampal microstimulation likely made the effect more powerful than a normally generated hippocampal output could have. On the other hand, the observed effects remained for several 100 ms after the termination of the activation pulses. In addition, a previous study showed that there can be a strong correlation between SI neurons in deep cortical layers and hippocampal neurons during oscillatory activity (Sirota et al., 2003). And, hippocampal neurons can be active at high firing rates (Tang et al., 2017; Viney et al., 2022), not seldom in relative synchrony (Harris et al., 2003). This suggests that hippocampal output can continually modify the global cortical state with high potency, including impacting the responses of SI neurons to tactile stimuli.

Note that the fact that the cluster analysis identified multiple preferred response states does not imply that these response states are fixed entities or the only response states possible (Etemadi et al., 2022; Norrlid et al., 2021). The cluster analysis merely indicates that there are some central response features that some responses share, whereas at least one of these features were not shared by the members of the other clusters. This is equivalent to that the cortical network provides a wide variety of network solutions that vary over time, and the moment the internal state is combined with a specific spatiotemporal input pattern, that network solution will unfold and result in a specific combination of response features. Importantly, the relative orthogonality between these response states, as indicated by the PCA/decoding analysis, indicate that the network in this regard may work according to attractor dynamics (Etemadi et al., 2022; Norrlid et al., 2021), conceptually explained in a recent review (Khona and Fiete, 2022). However, the present findings suggest that such attractor dynamics may be the product of the combined activity across very large areas of the central nervous system.

### Limitations of the study

It is inevitable that the anesthesia to some extent limited the range of possible spontaneous activity states. We do not think that this invalidates the main observed principle that the hippocampal perturbation tended to shift the state of the global network such that the range of network solutions to each given sensory input pattern would alter. This notion comes from an analysis of the multiple intracellular response states induced by the same TA input patterns used here (Norrlid et al., 2021), where we found that the response states arose both during the synchronized ECoG state, the probability of which is increased by the anesthesia, and during desynchronized activity, which is more similar to the predominant state observed in awake animals (Constantinople and Bruno, 2011). Furthermore, the physiological network structure, quantified as the recruitment order of a high number of simultaneously recorded cortical neurons across different sources of network excitation, does not alter with anesthesia (Luczak and Bartho, 2012; Luczak et al., 2009). Also, the type of network attractor dynamics that we observed here is likely closely related to the type of neuron population level dynamics observed in the hippocampus (Gardner et al., 2022), which remains stable whether the animal is awake or in deep sleep (Gardner et al., 2019).

### Broader implications of our findings

Our findings speak to the fundaments of the mode of operation of the cortical network. Several theories of neocortical circuitry function assume an ordered sequential hierarchical processing. But recent neuroanatomical studies indicate that the profusion of global interconnections across the neocortical neuronal networks has been vastly underestimated (Gerfen et al., 2018) (https://www.janelia.org/project-team/mouselight; https://mouse.braindatacenter.cn/) and that also the cortico-thalamo-cortical pathways are much more widespread (Halassa and Sherman, 2019) than what is typically accounted for in models of cortical network operation. Indeed, several recent analyses at the cellular level indicate that tactile information is globally distributed in the neocortex (Enander and Jorntell, 2019; Enander et al., 2019; Muret et al., 2022; Wahlbom et al., 2019; Wahlbom et al., 2021). Hence, these earlier findings speak in favor of the cortex being a highly recurrent network, rather than having a predominantly feed-forward network architecture. Here we show that this recurrent network seems to include even the hippocampus.

In a recurrent cortical neural network architecture, it is to be expected that a tactile input pattern, or in principle any other sensory input pattern, would combine with an ever evolving state of the neocortical network. This state would equal a component of the ‘‘prior,’’ ‘‘expectation,’’ or ‘‘prediction’’ currently residing in the neocortical network. That internal state component will be an important determinant of the resulting activity distribution across all neurons of the cortical circuitry that results when the tactile input pattern arrives—which is a potential definition of a ‘‘percept’’. Our results indicate that the cortical response states are profoundly impacted by hippocampal output. Whatever information that is being processed in the hippocampus will hence to some extent impact how we perceive a given tactile input, which for example could contribute to semantic and emotional aspects in haptic perception (Albert et al., 2016; Iosifyan et al., 2017; Rolls, 2022).

## Author Contributions

LE and HJ designed the experiments. LE performed the experiments. JMDE designed and performed the clustering analysis. LE, JMDE and HJ analyzed the results, and wrote the article.

## Acknowledgements

This work was supported by the EU H2020 Grant FETOpen project# 829186, ‘ph-coding’ (Predictive Haptic COding Devices In Next Generation interfaces)

## STAR Methods

### Resource availability

#### Lead contact

For any additional inquiry and information related to materials and resources used in this work Henrik Jörntell is the lead contact.

#### Materials availability

This study did not use any new/unique materials and/reagents.

#### Data and code availability

- The original contributions and raw data belong to this study will be deposited in figshare.com and made publicly available on acceptance
- The supporting data will be deposited at figshare.com and will be publicly available on acceptance
- Further information about the analyzed data discussed in this paper will be provided by the lead contact upon demand.

## Experimental model and subject details

We made recordings from 9 anesthetized male adult Sprague Dawley rats (weight: 250 – 380 g, age: 12 ± 2 weeks) in acute preparations. Before experiments, animals were maintained by Lund University animal facilities with 12h light/dark condition, 2-3 animals per cage (type 3H), and with free access to food and water.

### Institutional permission

Ethical approval for this research was obtained in advance from the local animal ethics committee in Lund/Malmö (Ethical permit ID: M13193-2017).

## Method details

### Surgical procedures

In order to make acute in vivo recordings, adult Sprague Dawley rats were initially prepared in the same way as in previous studies (Etemadi et al., 2022; Norrlid et al., 2021). Briefly: 1) The animal was sedated by inhaling air mixed with isoflurane gas (3%, for 2 min); 2) To induce general anesthesia, a mixture of ketamine/xylazine (ketamine: 40 mg/kg and xylazine: 4 mg/kg, accordingly) was injected intraperitoneally; 3) an incision in the inguinal area of the hindlimb was made to insert a catheter in the femoral vein for continuous infusion of Ringer acetate and glucose mixed with anesthetic (ketamine and xylazine in a 20:1 ratio, delivered at a rate of 5 mg/kg/h ketamine). 4) A hemicraniectomy (∼2 × 2 mm) was performed on the the right side of the skull at the area covering the somatosensory cortex (SI) on the right-hand side (∼2 × 2 mm), located at (from bregma): Ap: - 1.0 - +0.1, ML: 3.0–5.0.

A second cortical exposure over the visual cortex was made to gain access to the caudal CA1 and the subiculum complex of the hippocampus, located at (from bregma): AP: -6, ML: 3. The stimulation electrode was advanced 5.8 mm at an angle of 8 – 10 degrees to reach the subiculum complex. For the entire experiment, the EcoG was recorded via a surface electrode placed in the second cortical exposed area. To prevent and tissue dehydration, the second exposed cortical area was covered exposed cortical areas with liquid paraffin oil. To stabilize the tissue and to prevent dehydration, the first cortical exposure was covered in agarose.

The anesthetics used were chosen because they have previously been reported to not dramatically alter the neuronal recruitment order in spontaneous activity fluctuations and in stimulation-evoked responses as compared to the awake animal (Luczak and Bartho, 2012). As we have previously discussed extensively (Norrlid et al., 2021), anesthesia was required in order to achieve identical tactile stimulation patterns (where the electro tactile interface was the key, but would not be accepted in the awake animal) over a sufficiently long period of time. It also served to minimize brain activity noise caused by uncontrollable movements and internal thought processes unrelated to the stimuli. The level of anesthesia was assessed both by regularly verifying the absence of withdrawal reflexes to noxious pinch of the hind paw and by continuously monitoring the irregular presence of sleep spindles mixed with epochs of more desynchronized activity, a characteristic of sleep (Niedermeyer and da Silva, 2005). The animal was sacrificed by an overdose of pentobarbital at the end of the experiment. For histological evaluations, the animals were perfused with 4% paraformaldehyde (PFA) at the end of the experiment and the brain was removed for further processing (see below).

### Data collection: *In vivo* neuronal recordings

In order to analyze synaptic inputs intracellularly, we performed whole cell recordings in the current clamp mode. Patch pipettes were made from borosilicate glass capillaries pulled to 6 – 10 MΩ of impedance, using a Sutter Instruments P-97 horizontal puller. The pipettes were back-filled with an electrolyte solution of (in mM): potassium gluconate (135), HEPES (10), KCl (6.0), Mg-ATP (2), EGTA (10). The solution was titrated to pH of 7.35–7.40 using 1 M KOH. Recording signals were amplified using the EPC-800 patch-clamp amplifier (HEKA Elektronik) in the current-clamp mode (bandwidth from DC up to 100 kHz). Throughout recording sessions, time continuous data was acquired and digitized at 100 kHz using the CED 1401 mk2 hardware controlled from the Spike2 software (Cambridge Electronic Devices, CED, Cambridge, UK).

The patch electrode was inserted into the SI area (Paxinos and Watson, 2006) and advanced in a stepwise manner with applied positive pressure to prohibit blockage of the electrode tip. The pipette was advanced in a slow forward movement (⁓ 0.28 – 0.3 um/sec) until a sudden change in the electrode resistance was detected. When a dramatic increase in the magnitude of the isolated spike, evoked or spontaneous, occurred the positive pressure was replaced by a brief negative pressure. Also, to create a more optimal condition for the establishment of a GigaOhm seal, a mild hyperpolarizing current (in the order of -10 pA) was applied to the electrode. Then, to reach to the intracellular space episodical negative pressures were applied to the electrode tip. The data recording started after obtaining low noise access to the intracellular environment with a stable signal. The recordings were characterized by stable membrane potential of <-55 mV in down states, and with a peak-to-peak value between up and down states of >10 mV, with spike amplitudes of >25 mV in the beginning and end of stimulation protocols. All neurons recorded were putative pyramidal neurons rather than interneurons based on that they exhibited infrequent bursts of two or three spikes but had an absence of longer bursts or sustained periods of high firing (Luczak et al., 2009). All recordings were made in the neocortical layers II/III-V (350 – 1100 um of depth).

### Design: Electrical tactile and cortical stimulations

Once an intracellular recording was obtained, the experiments consisted of delivering tactile stimulation patterns through an electro tactile interface, combined with intracortical electrical perturbations. Four bipolar pairs of percutaneous stainless steel needle electrodes (isolated except for the tip (0.2-0.5 mm) were inserted into the superficial part of the skin of the second digit of the contralateral forepaw). Each pair represented a single channel for providing tactile stimulation (Figure 1A). Each electrode pair delivered a 400 µA constant current pulse with a duration of 200 µsec, which is about two times the threshold for activating tactile afferents using this type of stimulation (Bengtsson et al., 2013), and below the threshold for activating nociceptive afferents (Ekerot et al., 1987).

We used eight predefined spatiotemporal tactile afferent (TA) stimulation patterns (F5, S5, F10, S10, F20, S20, F∞, S∞; Fig. 1A). These patterns were delivered to the skin in a preset randomized order as previously reported (Enander et al. (2019); (Norrlid et al., 2021); Oddo et al. (2017); Wahlbom et al. (2019)). The duration of each TA stimulation pattern was less than 350 ms, and a randomized time interval of about 1.8 s separated the consecutive TA input patterns (Oddo et al., 2017).

Some of these TA inputs were combined with localized microstimulation in the subiculum/terminal part of the CA1 of the hippocampus. We used a glass-insulated tungsten electrode as previously described (Jorntell and Ekerot, 1999). In brief, the exposed tip of the electrode was about 100-150 um to deliver the stimulation against a grounding silver wire electrode inserted into the neck muscle. The microstimulation (HIP) consisted of constant current pulses of 0.2 ms duration and 0.4 mA intensity, delivered in trains of 7 pulses separated by 3 ms. When the HIP stimulation was combined with TA stimulation, the first TA stimulation pulse occurred with a 17 ms delay relative to the last pulse of the HIP stimulation train. This was done as we were not interested in the direct effects of HIP stimulation on the processing of the TA input, but rather the long lasting state change in the cortical network induced by HIP. Each of the 8 TA input patterns were repeated 50 times in isolation and 50 times in combination with the HIP stimulation (obtaining a total of 16 stimulation patterns) (only one of the 11 cells failed to reach 50 repetitions and instead had 44 repetitions of all stimulation patterns). All stimulation patterns were delivered in a pseudo-random order (i.e. a random order that was re-used across each experiment). In the cases where the first protocol was completed, we also applied the HIP stimulation in isolation (100 repetitions separated by 1.2 s) (N= 9 recordings). In some cases (N=7 recordings), where the neuron recording continued to be of a high quality, additional rounds of the protocol were commenced (for a total of 61 to 150 repetitions).

## Histological process, clearing technique

### Cresyl violet acetate staining

For postmortem assessment of the position of the stimulation electrode, animals were perfused with 4% PFA. At the end of each experiment, an i.v. overdose of ketamine/xylazine followed by transcardial perfusion with volume of ∼150 ml room temperature saline (0.9% NaCl in distilled water) and then 300ml of 4% paraformaldehyde (PFA) in 0.1 M phosphate buffer (pH 7.4), which was kept cold on the ice. The extracted brain post fixated in 4% PFA for 48h and then cryoprotected by immersion in a 25% sucrose solution. After obtaining the equilibrium, the brain was frozen on dry ice and kept in the freezer. Then, the 25 µm thick sections (cryostat, Microm GmbH, HM 560, Germany) mounted on glass (Super Frost^®^ plus slides, Mänzel-Gläser, Germany) were kept at -25 ^⸰^C in the freezer. For the staining process, mounted sections were immersed in ethanol/chloroform (1:1) overnight at room temperature (RT). In next day, the process followed by rehydration in ethanol (95%), rinsed in distilled water (2 min./step), stained in Cresyl violet (Life Science product & service company; 0.1% in 0.3% acetic acid in dH_2_O; 5 min.), rinsed in distilled water until sections became clear of extra dye, followed by dehydration in ethanol 95% (2 times 2 mins), and clearing in xylene (2×100%, 5 min.). At the end, the cover slip was placed on the stained sections using the DPX mounting media (Fluka, Germany). Slides were examined under light microscopy and images were obtained using DS-Ri1 digital camera (Nikon Instruments, Japan) mounted on a Nikon Eclipse 80i microscope.

## Immunolabeling and iDISCO Clearing technique for light sheet microscopy

### Sample Pretreatment with Methanol

For some brains we used light sheet microscopy to visualize the positions of the SI recording electrode and the HIP stimulation electrode. We used a standard iDISCO protocol (idisco.info) for PFA fixated frozen brain sample (one sample/ one hemisphere). iDISCO is a method recognized for its ability in visualizing voluminous structures, therefore, the cellular profiles could be immunolabeled while retaining their morphological and molecular properties (Renier et al., 2014). In short, to de-freeze samples, they were washed in PBS (2x, 30 min. each). Then, they went through a dehydration procedure, in different series of methanol/PBS dilutions: 20%, 40%, 60%, 80%, and in methanol 100% (2x)1 hr/step. Then, they were incubated overnight in 66% DCM/ 33% methanol, RT with slight shaking (60 RPM) using the Ultra Rocker (Bio-Rad Laboratories, Sweden) in100% methanol (2x, 30 min each), then remained 1h at 4 ^⸰^C. Samples were then bleached in chilled fresh 5% H2O2 in methanol (1 volume 30% H2O2 to 5 volumes of methanol, ice cold) over the night at 4 ^⸰^C. Next day, the brain was rehydrated with methanol/PBS series dilutions: 80%, 60%, 40%, 20%, PBS, 1 hr/step, RT.

### Immunolabeling Protocol

Neurobiotin (Vector Laboratories, SP-1120) was included in the *in vivo* patch-clamp recording electrode solution (1.5% dilution). We also coated the HIP stimulation electrode in Neurobiotin for assessing its location within the brain. To increase the chances of detecting the tracer in the light sheet microscopy, we used a primary antibody against neurobiotin. Hence, bleached samples were incubated in the permeabilization solution buffer (PTx.2/glycine/DMSO) at 37 ^⸰^C for 5 days (d), then blocked in PTx.2/DMSO/Donkey Serum (Jackson laboratories) at 37 ^⸰^C for 5 d. Then they were incubated with the primary antibody (neurobiotin) (1:1000) in PTwH (PBS/0.2% Tween-20 with 10 µg/ml heparin)/5% DMSO/3% Donkey Serum for 9 days. Then, they were washed with PTwH for 10, 15, 20 min, 1 hr, and 4-5 times more until the next day, followed by a 10 day incubation with the secondary antibody, streptavidin Alexa Flour-568 conjugated (1:500) (ThermoFisher, Catalog number: S11226) in PTwH/3% Donkey Serum at 37 ^⸰^C. Samples were finally washed in PTwH for 10, 15, 20 min, 1 hr and then 4-5 times before the clearing step the next day. For all these steps samples were under slight shaking condition.

### Tissue Clearing

Immunolabeled samples were cleared following iDISCO method (idisco.info). Samples were dehydrated in methanol/PBS series: 20%, 40%, 60%, 80%, 1h each, twice with 100% methanol 1hr each at RT and left over the night. Next day, they were incubated in 66% DCM/33% methanol, 3 hr, RT, shaking. Then, incubated in 100% DCM (Sigma 270997-12X100ML), 15 min twice, shaking. In the final step they were incubated in Ethyl Cinemate (Sigma-Aldrich, SKU: 112372-100G) until clear and stored in RT until imaging with light sheet microscopy.

### Light-Sheet Microscopy

The cleared hemisphere imaged with an Ultra Microscope II (LaVision Biotec) carrying an sCMOS camera (Andor Neo model 5.5-CL3), which the objective lens 1.3X (LaVision LVMI-Fluor 1.3×70.08 MI Plan) was used and 600/30 emission for Alexa Fluor 568. Stacks were made by overlapping up to 10% and stitched via Arivis Vision 4D 3.01 software.

## Quantification and statistical analysis

### Post-processing of recorded neuronal data

The raw neuronal membrane voltage recordings were post-processed by first defining a stable baseline voltage by applying a least-square linear detrending to 10 second long segments. The recordings were subsequently low pass filtered using a rolling average of 100 bins equating to a boxcar window with duration of 1 ms. In recordings where occasional spikes appeared, the onset of these spikes were found using a spike shape template and a recursive fitting algorithm (Mogensen et al., 2019). Stimulation artefacts and any spurious action potentials were removed by linear blanking, with a duration verified to be sufficient to blank out the events by visual inspection. Finally, the recordings were down sampled to 1 kHz. Overall, the software used in the below analysis was NumPy and SciPy, standard Python libraries in data science.

### Definition of evoked responses

The time window included in the analysis related to response clustering below, the analysis included full raw data sweeps starting from 5 ms after the onset of the TA stimulation (i.e., for both TA and HippTA stimulation patterns) and ending 400 ms later (some TA patterns lasted almost 350 ms, and responses could sometimes be detected at least 50 ms after the last stimulation pulse). The 5 ms gap relative to the TA onset corresponded to the conduction time from the skin stimulation to the earliest possible arrival of synaptic responses in the neocortex.

### Clustering method

We used the same method to cluster the responses as we used in a previous publication (Etemadi 2022). This identifies clusters of similar responses, with respect to their time-voltage curves, evoked by the same stimulation pattern, in an unsupervised manner. Hence, the algorithm was designed to identify clusters where the ‘members’ of each group were internally similar, while also being distinct relative to the other responses evoked by the same stimulation pattern. Furthermore, the algorithm had to be able to automatically determine the number of clusters that could exist, since this could not be known *a priori*.

First, each response was z-score normalized, i.e., the response mean over the whole time period was subtracted from each response and then the remainder of each response was divided by the standard deviation. Secondly, a distance matrix was calculated for the whole set of responses. The distance matrix was populated by calculating the M number of Principal Components (PC) explaining 95% of the variance for N-1 responses. All responses were then fitted to the PCs using the least square method. The resulting PC coefficients were used to position the responses in the M-dimensional PC space. Within this M-dimensional space, the Euclidean distances between a reference response and the rest of responses were calculated. The distances between the reference response and each of the other responses were then put in a matrix and the procedure was repeated until each response had been the reference. Finally, the distances were normalized to a 0-1 range.

From the Euclidean distance calculations above, a linkage matrix was calculated using the hierarchical clustering approach of Ward’s minimum variance method (Ward, 1963). This method sequentially finds the two closest neighboring responses with respect to their Euclidean distance. Then it finds the pair of the second closest neighboring responses and so on until there are no responses left. However, a response may also be closer to a center point between a pair of responses than it is to other responses, in which case the distance to that center point is assigned to the response. This distance is called the cophenetic distance. Sometimes, the shortest distance that can be found is between two such center points. All these distances are used to create the hierarchical structure of responses, where the distances between all such pairs can be compared.

The next step is to identify the cophenetic distance at which the identified clusters are objectively the most separable. The total cophenetic distance of a hierarchy was divided into 100 steps. Each level of cophenetic distance was used it as a threshold, which would separate clusters from each other (i.e. a cluster of responses with a cophenetic distance larger than the threshold was considered a unique cluster), whereby an assignment of a cluster id was made. At this point we could perform data shuffling if needed. Data shuffling was achieved by shuffling the leaf position of each response in the hierarchy (The shuffling was performed using the NumPy shuffle method), therefore disrupting the relationship between cluster and response.

We next calculated the separability of the clusters by using a decoding analysis (described in detail in ‘Evaluation of cluster separability using decoding analysis’ below). The decoding analysis yielded, for each clustering threshold, a measure of the separability of the clusters as the F1-score. With the F1-score, we could apply Gap statistics to identify the cophenetic distance providing the largest cluster separation relatively to the shuffled data. Meaning that for each of the 100 cophenetic distances considered, we obtained a F1 score for the non-shuffled data and another F1 score for the shuffled data, subtracting the F1-score of the shuffled data from the F1-score of the actual data yielded the difference also known as the Gap Statistic (Tibshirani et al., 2001). The calculation of the Gap Statistic was repeated three times per clustering threshold and the mean value was reported. Since the specific value of the chosen cophenetic distance defines the number of clusters to be considered, we could identify continuous stretches of cophenetic distances where the number of clusters, and the clusters themselves, remained unchanged. Over each of these continuous stretches we calculated a mean Gap Statistic. The middle value of cophenetic distance of the stretch with the maximum Gap statistic was used to define the cophenetic distance at which the clustering was objectively the most distinct. The clustering obtained at this value was then used for the remainder of the analysis and displays.

While following the above principles, there were some special situations that could arise. A cluster was assumed to be valid only if it had five or more members. All non-valid clusters were put in the same “undefined” cluster. In addition, if the situation arose that one of the clusters consisted of only one member this response and cluster was excluded from all subsequent analysis and displays.

### Evaluation of cluster separability using decoding analysis

We used the same general decoding algorithm to determine the specificity of the response clusters as in several previous publications (Enander and Jorntell, 2019; Enander et al., 2019; Wahlbom et al., 2019; Wahlbom et al., 2021), including analysis of intracellular recording data with multiple response states to each tactile input pattern (Norrlid et al., 2021).

The responses and their respective label were split into a training- and a test set (80%/20% split) stratified by cluster label frequency. The training set was used to train a PCA model explaining 95% of the variance, and the coefficients obtained by fitting the responses with the least-square-method to the principal components were calculated for both the training set and the test set. Finally, a kNN-classification was performed with the training set coefficients as training data, and the test set coefficients as test data. This decoding algorithm was performed 30 times, each time with a new split of train and test data. The F1-score was calculated from the average classification result across the 30 repetitions.

The decoding analysis was also repeated with the randomly shuffled cluster labels (as described in Clustering method), reported as the “shuffled” context. This was done to establish the actual chance level of the data, as opposed to the theoretical chance decoding level (which equals 1 divided by the number of clusters).

## REFERENCES

Ahrlund-Richter, S., Xuan, Y., van Lunteren, J.A., Kim, H., Ortiz, C., Pollak Dorocic, I., Meletis, K., and Carlen, M. (2019). A whole-brain atlas of monosynaptic input targeting four different cell types in the medial prefrontal cortex of the mouse. Nat Neurosci 22, 657–668.

Albert, B., De Bertrand De Beuvron, F., Zanni-Merk, C., Maire, J.-L., Pillet, M., Charrier, J., and Knecht, C. (2016). A smart system for haptic quality control: A knowledge-based approach to formalize the sense of touch. In International Joint Conference on Knowledge Discovery, Knowledge Engineering, and Knowledge Management (Springer), pp. 173–190.

Bengtsson, F., Brasselet, R., Johansson, R.S., Arleo, A., and Jorntell, H. (2013). Integration of sensory quanta in cuneate nucleus neurons in vivo. PLoS One 8, e56630.

Bullmore, E., and Sporns, O. (2009). Complex brain networks: graph theoretical analysis of structural and functional systems. Nat Rev Neurosci 10, 186–198.

Constantinople, C.M., and Bruno, R.M. (2011). Effects and mechanisms of wakefulness on local cortical networks. Neuron 69, 1061–1068.

Ekerot, C.F., Gustavsson, P., Oscarsson, O., and Schouenborg, J. (1987). Climbing fibres projecting to cat cerebellar anterior lobe activated by cutaneous A and C fibres. Journal of Physiology (London) 386, 529–538.

Enander, J.M.D., and Jorntell, H. (2019). Somatosensory Cortical Neurons Decode Tactile Input Patterns and Location from Both Dominant and Non-dominant Digits. Cell Rep 26, 3551–3560 e3554.

Enander, J.M.D., Spanne, A., Mazzoni, A., Bengtsson, F., Oddo, C.M., and Jorntell, H. (2019). Ubiquitous Neocortical Decoding of Tactile Input Patterns. Front Cell Neurosci 13, 140.

Etemadi, L., Enander, J.M.D., and Jorntell, H. (2022). Remote cortical perturbation dynamically changes the network solutions to given tactile inputs in neocortical neurons. iScience 25, 103557.

Gardner, R.J., Hermansen, E., Pachitariu, M., Burak, Y., Baas, N.A., Dunn, B.A., Moser, M.B., and Moser, E.I. (2022). Toroidal topology of population activity in grid cells. Nature 602, 123–128.

Gardner, R.J., Lu, L., Wernle, T., Moser, M.B., and Moser, E.I. (2019). Correlation structure of grid cells is preserved during sleep. Nat Neurosci 22, 598–608.

Gerfen, C.R., Economo, M.N., and Chandrashekar, J. (2018). Long distance projections of cortical pyramidal neurons. J Neurosci Res 96, 1467–1475.

Halassa, M.M., and Sherman, S.M. (2019). Thalamocortical Circuit Motifs: A General Framework. Neuron 103, 762–770.

Harris, K.D., Csicsvari, J., Hirase, H., Dragoi, G., and Buzsaki, G. (2003). Organization of cell assemblies in the hippocampus. Nature 424, 552–556.

Iosifyan, M., Korolkova, O., and Vlasov, I. (2017). Emotional and semantic associations between cinematographic aesthetics and haptic perception. Multisensory Research 30, 783–798.

Jorntell, H., and Ekerot, C.F. (1999). Topographical organization of projections to cat motor cortex from nucleus interpositus anterior and forelimb skin. J Physiol 514 (Pt 2), 551–566.

Khona, M., and Fiete, I.R. (2022). Attractor and integrator networks in the brain. Nat Rev Neurosci 23, 744–766.

Luczak, A., and Bartho, P. (2012). Consistent sequential activity across diverse forms of UP states under ketamine anesthesia. Eur J Neurosci 36, 2830–2838.

Luczak, A., Bartho, P., and Harris, K.D. (2009). Spontaneous events outline the realm of possible sensory responses in neocortical populations. Neuron 62, 413–425.

Matsumoto, N., Kitanishi, T., and Mizuseki, K. (2019). The subiculum: Unique hippocampal hub and more. Neurosci Res 143, 1–12.

Mogensen, H., Norrlid, J., Enander, J.M.D., Wahlbom, A., and Jorntell, H. (2019). Absence of Repetitive Correlation Patterns Between Pairs of Adjacent Neocortical Neurons in vivo. Front Neural Circuits 13, 48.

Muret, D., Root, V., Kieliba, P., Clode, D., and Makin, T.R. (2022). Beyond body maps: Information content of specific body parts is distributed across the somatosensory homunculus. Cell Rep 38, 110523.

Niedermeyer, E., and da Silva, F.H.L. (2005). Electroencephalography: basic principles, clinical applications, and related fields (Lippincott Williams & Wilkins).

Norrlid, J., Enander, J.M.D., Mogensen, H., and Jorntell, H. (2021). Multi-structure Cortical States Deduced From Intracellular Representations of Fixed Tactile Input Patterns. Front Cell Neurosci 15, 677568.

Oddo, C.M., Mazzoni, A., Spanne, A., Enander, J.M., Mogensen, H., Bengtsson, F., Camboni, D., Micera, S., and Jorntell, H. (2017). Artificial spatiotemporal touch inputs reveal complementary decoding in neocortical neurons. Sci Rep 8, 45898.

Paxinos, G., and Watson, C. (2006). The rat brain in stereotaxic coordinates: hard cover edition (Elsevier).

Pereira, A., Ribeiro, S., Wiest, M., Moore, L.C., Pantoja, J., Lin, S.C., and Nicolelis, M.A. (2007). Processing of tactile information by the hippocampus. Proc Natl Acad Sci U S A 104, 18286–18291.

Renier, N., Wu, Z., Simon, D.J., Yang, J., Ariel, P., and Tessier-Lavigne, M. (2014). iDISCO: a simple, rapid method to immunolabel large tissue samples for volume imaging. Cell 159, 896–910.

Ringach, D.L. (2009). Spontaneous and driven cortical activity: implications for computation. Curr Opin Neurobiol 19, 439–444.

Rolls, E.T. (2022). The hippocampus, ventromedial prefrontal cortex, and episodic and semantic memory. Progress in Neurobiology, 102334.

Schultz, C., and Engelhardt, M. (2014). Anatomy of the hippocampal formation. Front Neurol Neurosci 34, 6–17.

Sirota, A., Csicsvari, J., Buhl, D., and Buzsaki, G. (2003). Communication between neocortex and hippocampus during sleep in rodents. Proc Natl Acad Sci U S A 100, 2065–2069.

Tang, W., Shin, J.D., Frank, L.M., and Jadhav, S.P. (2017). Hippocampal-Prefrontal Reactivation during Learning Is Stronger in Awake Compared with Sleep States. J Neurosci 37, 11789–11805.

Tibshirani, R., Walther, G., and Hastie, T. (2001). Estimating the number of clusters in a data set via the gap statistic. Journal of the Royal Statistical Society: Series B (Statistical Methodology) 63, 411–423.

Viney, T.J., Sarkany, B., Ozdemir, A.T., Hartwich, K., Schweimer, J., Bannerman, D., and Somogyi, P. (2022). Spread of pathological human Tau from neurons to oligodendrocytes and loss of high-firing pyramidal neurons in aging mice. Cell Rep 41, 111646.

Wahlbom, A., Enander, J.M.D., Bengtsson, F., and Jorntell, H. (2019). Focal neocortical lesions impair distant neuronal information processing. J Physiol 597, 4357–4371.

Wahlbom, A., Enander, J.M.D., and Jorntell, H. (2021). Widespread Decoding of Tactile Input Patterns Among Thalamic Neurons. Front Syst Neurosci 15, 640085.

Ward, J.H. (1963). Hierarchical grouping to optimize an objective function. Journal of the American statistical association 58, 236–244.

Winnubst, J., Bas, E., Ferreira, T.A., Wu, Z., Economo, M.N., Edson, P., Arthur, B.J., Bruns, C., Rokicki, K., Schauder, D., et al. (2019). Reconstruction of 1,000 Projection Neurons Reveals New Cell Types and Organization of Long-Range Connectivity in the Mouse Brain. Cell 179, 268–281 e213.

